# Design and fabrication of demountable 3D microphysiological systems for modeling barrier function and underlying tissue interactions

**DOI:** 10.1101/2025.10.10.681761

**Authors:** Selina Banerjee, Ryan Brady, Lina Abu-Absi, Dylan Miller, Bryan Schellberg, Guohao Dai, Abigail Koppes, Ryan Koppes

## Abstract

Several recent advances in microphysiological systems (MPSs) or organ-on-chip technology have demonstrated its potential for replacing traditional *in vitro* and animal models in the coming years. Despite the physiological relevance and cost-effectiveness of organ chips, there are several hurdles that must be overcome for widespread adoption for biological studies. Many shortcomings of manufacturing and scalability have been overcome by a transition from PDMS to thermoplastics. However, challenges have arisen in these sealed, brittle systems related to end-point tissue analyses, harvest, and high-resolution imaging, which is particularly difficult for multi-layer organ chips. Here, we present low-cost organ chips that are fluidically sealed but demountable, fabricated using a cut-and-assemble method without the need for cleanroom technologies. We have validated the capabilities of this method by demonstrating the culture of human aortic smooth muscle cells and induced pluripotent stem cell-derived neural cells, encapsulated in gelatin methacryloyl (GelMA) hydrogel on chip, for up to 27 days. The 3D culture layer of the organ chip was removed, and high-resolution images were obtained via immunostaining. Furthermore, these organ chips facilitate rapid redesign and manufacture for alternative tissue and/or interface systems. To our knowledge, this is the first innervated organ chip with multiple removable cell culture layers, as well as the first humanized nerve-artery model that includes a three-dimensional hydrogel culture. In future work, these unique features of our platform can be utilized for investigating the crosstalk mechanisms between different cell types in co-culture.

**Impact Statement:** We present here a new method for fabricating low-cost demountable organ-on-a-chip platforms. This method leverages our recent cut & assemble method for layered 3D organ chips comprised of gas impermeable thermoplastics.

## Introduction

In recent years, microphysiological systems (MPSs) have gained popularity due to their increased physiological relevance and tunability over traditional two-dimensional cell culture methods and animal models^1,2^. Despite extensive development at the bench, organ chips have yet to be robustly adopted. Many recent designs have pivoted away from polydimethylsiloxane (PDMS) to brittle thermoplastics^3-6^. PDMS absorbs hydrophobic molecules and is highly gas permeable^7-9^, which reduces its applicability for drug screening or studying gas exchange, such as investigations on hypoxic environments or reactive oxygen species. Our MPS comprised of polymethyl methacrylate (PMMA) and glass can facilitate tightly controlled investigations of volatile hydrogen sulfide in gut-chip models^10^.

The transition to more brittle materials however has introduced one major challenge: the inability to directly access the inner culture chambers for imaging or biological assays after initial seeding^11^. Though complex geometries and interfacing cell culture layers provide added physiological relevance, optical, proteomic, and genetic analysis of individual cellular components become a challenge. While some analysis techniques can be carried out via extraction of cells, this fails to preserve their spatial resolution and often results in considerable loss of sample resulting in poor yield, especially in permanently sealed systems. Tissue constructs that are three-dimensional in the z-axis possess limitations in imaging resolution, especially for multi-layer organ chips or organ chips with hydrogel-encapsulated cell cultures. A potential workaround, microphysiological systems (MPS) fabricated with removable components, which either utilize reversibly bonded components or clamping strategies allow access to internal components at the experimental terminus^11,12^. James Hickman and Michael Shuler along with the Hesperos Inc. team have highlighted the utility of demountable and reconfigurable systems across an extensive library of elegant designs and utilities^13,14^. However, these systems rely on soft lithography and PDMS gaskets. Shah et al. has reported the use of silicone rubber gaskets in an organ-chip model of gut epithelial monolayers and microbiome in anaerobic conditions^15^. Towards more biomimetic models, interfacing 3D tissues will be required, and disassembly of models would ease terminal tissue harvesting, spatial genetic or proteomic analysis, and high-resolution imaging.

Here, we present a novel method for 3D MPS platforms fabricated via our well-established laser cut and assemble method (Fig. 1)^3,4,6,10,16^ and assembled with rubber gaskets and a closure force. This process was implemented to develop a neurovascular organ chip with interfacing cell culture layers that are fluidically sealed but demountable. The organ chip uses a rapid cut-and-assemble method previously established by our lab to fabricate laser cut polymethyl methacrylate and polyethylene terephthalate layers as well as stereolithographic 3D printed clear resin caps^3,17^. After evaluating several elastomer materials based on cost, gas permeability, ability to be laser cut, and biocompatibility, we determined that Viton is most suitable for use as gaskets in our chip (Fig. 2, Table 1). This organ chip uses standard hardware to fluidically seal the various layers, which allows straightforward rearrangement of layers for different experimental configurations and demounting. Each organ chip costs ∼$8-11 and can be easily manufactured using widely available makerspace equipment. This system has been validated with a standard gut monolayer model as well as a more complex human neurovascular co-cultures to create a model of the physiological nerve-artery interface. We demonstrated the utility of this design by immunostaining for neuronal catalysts, which have transient expression in neurons and therefore often require high resolution imaging for visualization. Additionally, we demonstrate the potential of our organ chip for multi-channel imaging or multiplexed imaging experiments, where we stained and imaged the 3D (aortic smooth muscle cell culture) and 2D (human aortic endothelial cell culture) layers of the organ chip independently.

**Table 1.** Materials Selection Criteria for Elastomer Layer.

| Elastomer | Laser Cutting Suitability | Color | CO<sub>2</sub> Permeability ((cm<sup>3</sup>·mm/m<sup>2</sup>·day·atm)) | F11 Cell Viability |
| --- | --- | --- | --- | --- |
| Silicone | Poor | Semi-Transparent | 212000 | 89% |
| EPDM | Moderate | Black | 7693 | 50% |
| Neoprene | Moderate | White | 1940 | 92% |
| **Viton** | **Excellent** | **Black** | **508** | **89%** |

**Figure 1.**
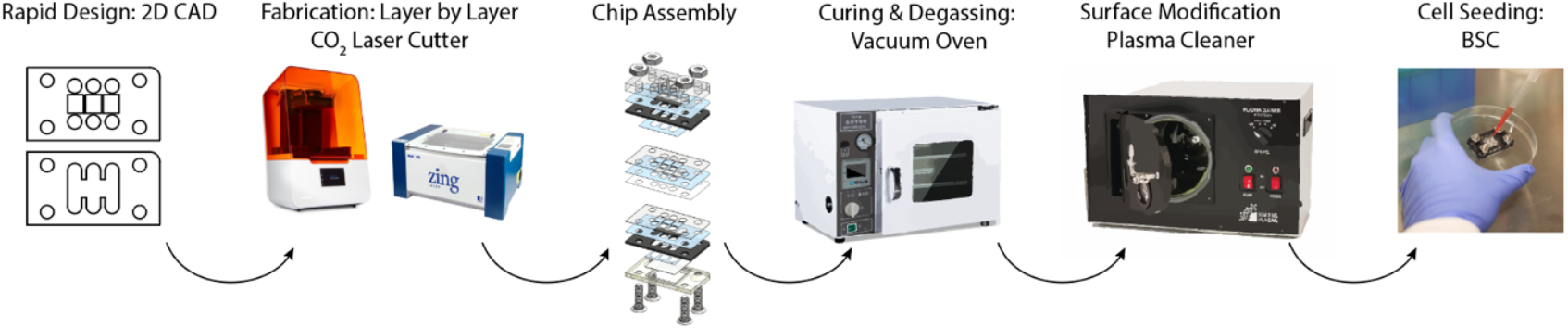
Laser cut & assemble method for fabricating demountable 3D MPSs. After assembly, MPSs were degassed and plasma treated before seeding of cells for co-culture experiments.

**Figure 2.**
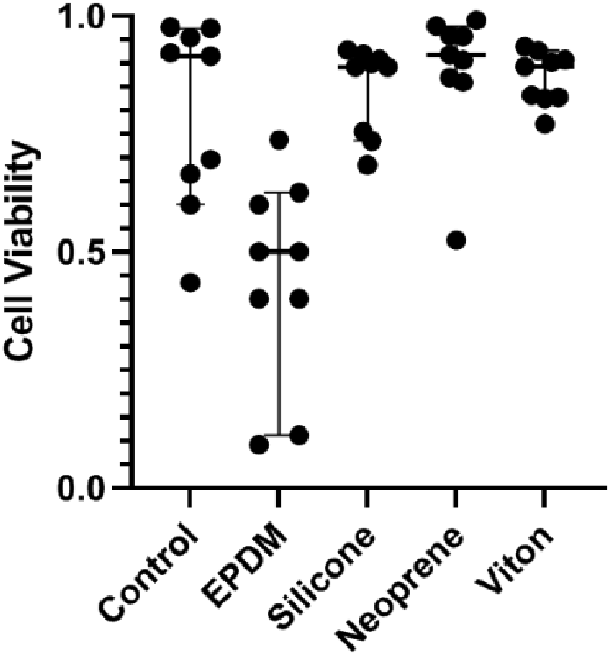
F11 cell viability cultured with elastomer samples for material selection.

## Materials and Methods

### Cell Viability and Material Selection

F11 (rat dorsal root ganglia – mouse neuroblastoma hybrid) cells (Millipore Sigma #08062601-1VL) were grown in basic media composed of Advanced DMEM (cat: 12491023, Gibco), fetal bovine serum (10%, Atlanta Biologicals), penicillin-streptomycin (1%, cat: 15140122, Gibco), and GlutaMAX (1X, cat: cat: 35050061, Gibco) between passage 26 and 40. Commercially available elastomers were evaluated for utility as an inert gasket layer (Table 1). Rubber samples were ordered in a sample pack (8450K11) including Silicone, Ethylene propylene diene monomer (EPDM), Neoprene, and Viton (1235N21) from McMaster Carr and cut into uniform square samples (10 by 10 mm). F11 cells were cultured in TC-treated 24 well plates at a density of 0.5 x 10^6^ cells/well with square samples for 7 days. Cells were then evaluated with a Live/Dead assay (ThermoFisher R37601), imaged, and quantified (Fig. 2).

### Device Fabrication

Neurovascular organ chip devices (Fig. 3) were constructed by a previously described cut-and-assemble method, with modifications (Fig. 1)^3,17^. Using a laser engraver system (Epilog Zing 16, Epilog Laser), double-sided adhesive tape (3M 966), 3/16” polymethyl methacrylate (PMMA, McMaster-Carr 8560K211), 0.0005” polyethylene terephthalate (PET, McMaster-Carr 8567K102), and 1/64” Viton were cut to the shapes of the respective layers as shown in Figure 4. Each layer of the organ chip device was assembled as follows:

**Figure 3.**
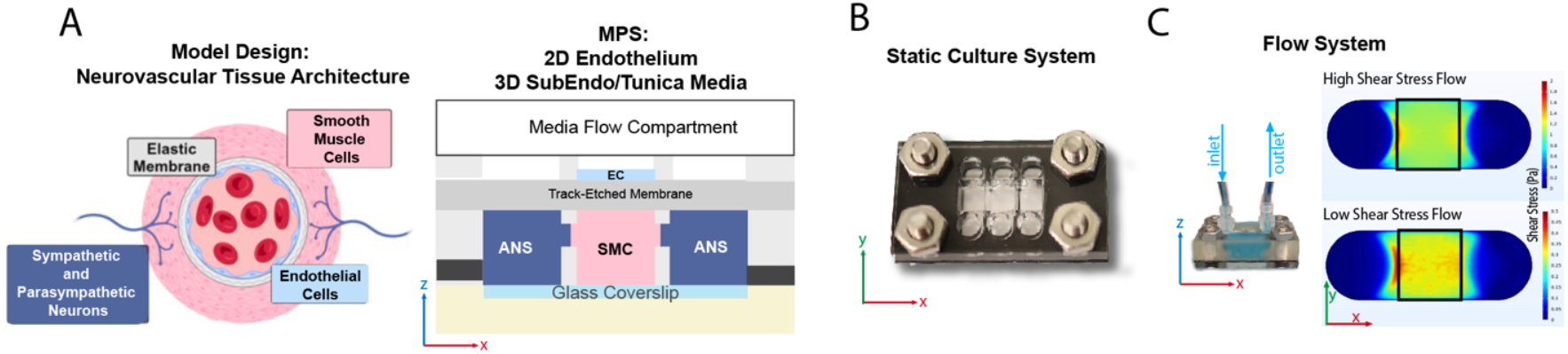
Demountable neurovascular MPS. A) Arterial structure, specifically the endothelium and tunica media, is recapitulated into simplified 3-chamber system. An endothelial monolayer is cultured on track etched membrane under a removable top to facilitate seeding, culturing, and fluidic flow. Endothelial monolayer directly interfaces with a 3D smooth muscle tissue representative of the tunica media. Abutting chambers contain adrenergic motor neurons representative of post ganglionic innervation. B) Top view of MPS C) Interchangeable top enables application of flow regimes to mimic high and low shear stress.

**Figure 4.**
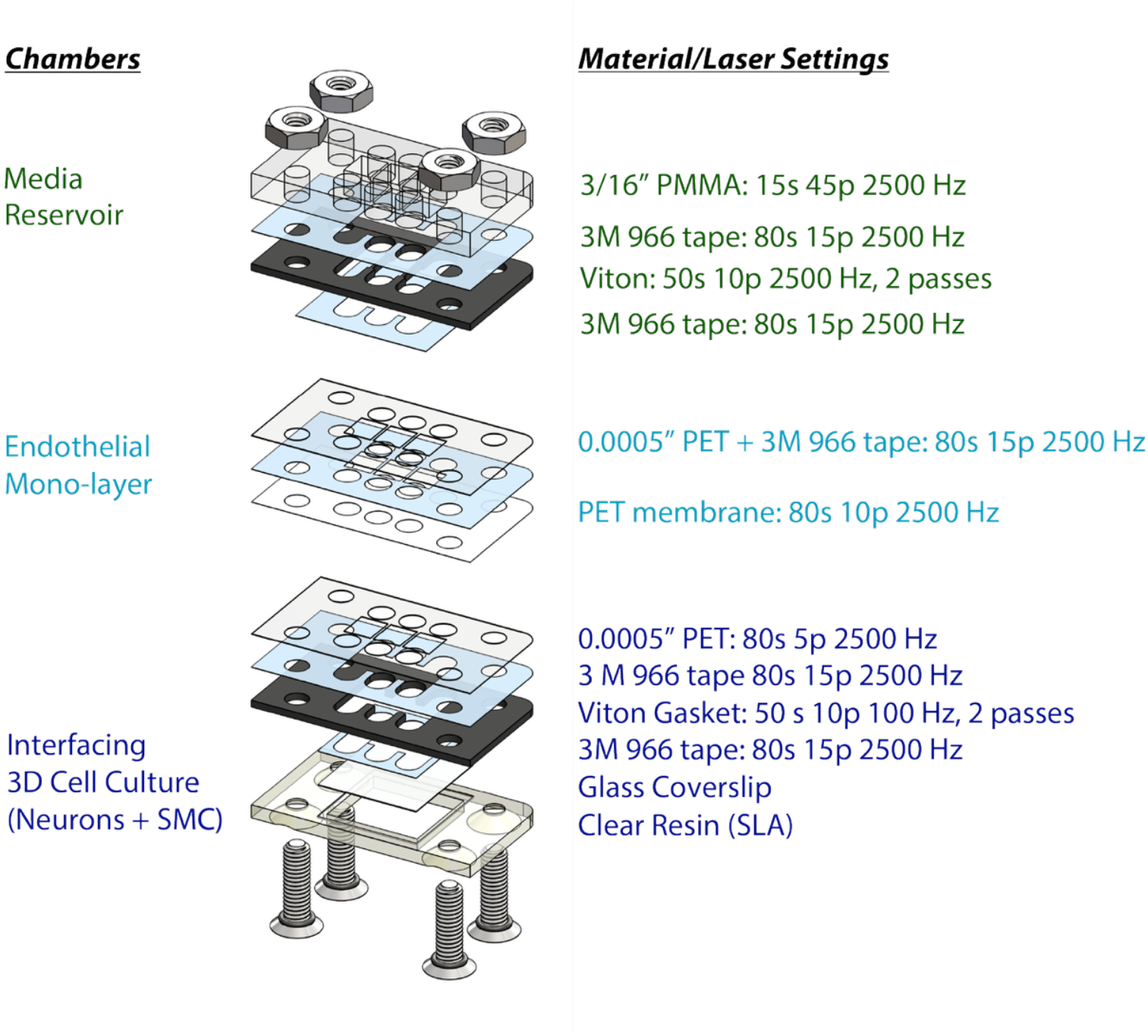
Layer by layer design of neurovascular chip and laser settings for cutting the different polymer materials throughout the chip assembly.

*To establish a media reservoir*, a Viton gasket piece was attached to a 3/16” PMMA piece with a piece of double-sided tape. This assembly was heat pressed at 53°C for ∼15 seconds. Then, a partial piece of double-sided tape was attached to the exposed side of the Viton gasket.

*To establish a 2D cell culture layer*, a double-sided tape sheet was attached to a sheet of 0.0005” PET on one side. The assembly was carefully attached to a PET membrane with 1 µm pore size (it4ip, 2000M12/620M103). Then, the media ports of the assembly were laser cut (Fig. 4). The assembly was then heat pressed at 53°C for ∼15 seconds. The partial piece of double-sided tape (which was previously attached to the media reservoir) was used to attach the 2D cell culture layer. Then, the media reservoir + 2D cell culture layer assembly was heat pressed again at 53°C for ∼15 seconds.

*To establish a 3D cell culture layer*, a 0.0005” PET piece (Fig. 4) piece was laser cut. A Viton gasket piece was attached to this PET piece with a double-sided tape, cut to the same shape as the Viton gasket. The assembly was then heat pressed at 53°C. A partial double-sided tape piece, which includes an 18 mm x 18 mm portion and a border to be detached, was attached to a #1 thickness 18 mm x 18 mm coverslip. For attachment of the glass coverslip (Fisherfinest 12548A), double sided adhesive tape was cut to an 18 mm x 18 mm square with borders. The coverslip attached to the center of the 18 mm x 18 mm tape, and the borders of the tape were then removed and discarded. The coverslip holder piece was fabricated by a stereolithographic 3D printer (Form 3B+, Formlabs) using Clear Resin (Formlabs). The coverslip with tape was then attached to the gasket on the assembly, and the assembly was placed on the stereolithographic (SLA) 3D printed clear resin coverslip holder. The assembly was heat pressed at 53°C for ∼15 seconds.

The three layers of the organ chip were assembled with 5/8” stainless steel screws with o-rings (McMaster Carr 98070A162) and nuts (McMaster Carr 91841A009). Chips were tightened to 0.12 Nm with an Anpuds digital Torque Wrench (3Nm) to ensure proper sealing, but not over tightening/compression. Assembled organ chips were vacuum incubated at 53°C for at least 96 hours. Then, organ chips were UV sterilized and plasma treated (Harrick Plasma PDC-001) for 180 seconds.

### Human iPSC Cultures and Differentiation Towards Autonomic Neuron Populations

Human induced pluripotent stem cells (iPSC, iPS(IMR90)-4) were sourced from WiCell. Cells were maintained at standard cell culture conditions (5% CO_2_, 37°C) in mTeSR Plus medium (STEMCELL Technologies 100-0276) on 6 well plates (Corning 07-200-83) coated with 1:100 Matrigel (Corning CB-40230) in DMEM-F12 (Gibco 11-320-033). Full media exchanges were performed daily. Accutase (Corning 25058CI) was used to disassociate cells when they approached confluency. ROCK inhibitor (Sigma Y0503) was added to cell culture media (10 μM) for ∼24 hours after thawing, before passaging, and after passaging.

Human iPSCs were differentiated following an adapted version of a protocol previously published by Takayama et al. [9] (Fig. 5). Briefly, iPSC at P18 + 42 were plated (Day -1) into a 6-well plate coated with 1:100 growth factor reduced Matrigel (Corning CB-40230) in DMEM-F12 (Gibco 11-320-033) at x 10^5^ cells/well and maintained at standard culture conditions (5 % CO_2_, 37°C) in mTeSR Plus medium (STEMCELL Technologies 100-0276) and 10 μM ROCK inhibitor (Sigma Y0503). The following day (Day 0), the ROCK inhibitor media was removed. hiPSCs were then differentiated with three separate base media compositions: a knockout serum replacement-based medium (KOSR), an N2 supplement based medium (N2), and a Neurobasal-A/B27+ based medium (B27). The KOSR based medium was composed of DMEM-F12 (Gibco 11-320-033), 20% KOSR (Gibco 10828010), 1% non-essential amino acids (Gibco 11140050), 0.5 mM monothioglycerol (Spectrum Chemical M11771), and 1% penicillin-streptomycin (Gibco 15140122). The N2 based medium was composed of DMEM-F12, 1% N2 supplement (Gibco 17502048), 1% non-essential amino acids, and 1% penicillin-streptomycin. Lastly, the B27 medium was composed of 50X B27 Plus Supplement (Gibco A3582801) diluted to 1X in Neurobasal-A (Gibco 10888022). The differentiation protocol was initiated on Day 0, and full media exchanges were completed every other day until Day 13, at which the frequency was every third day. On Day 16, medium was replaced for both sympathetic-like and parasympathetic-like cultures with B27 medium and both cell types were maintained in B27 medium until culture termination.

**Figure 5.**
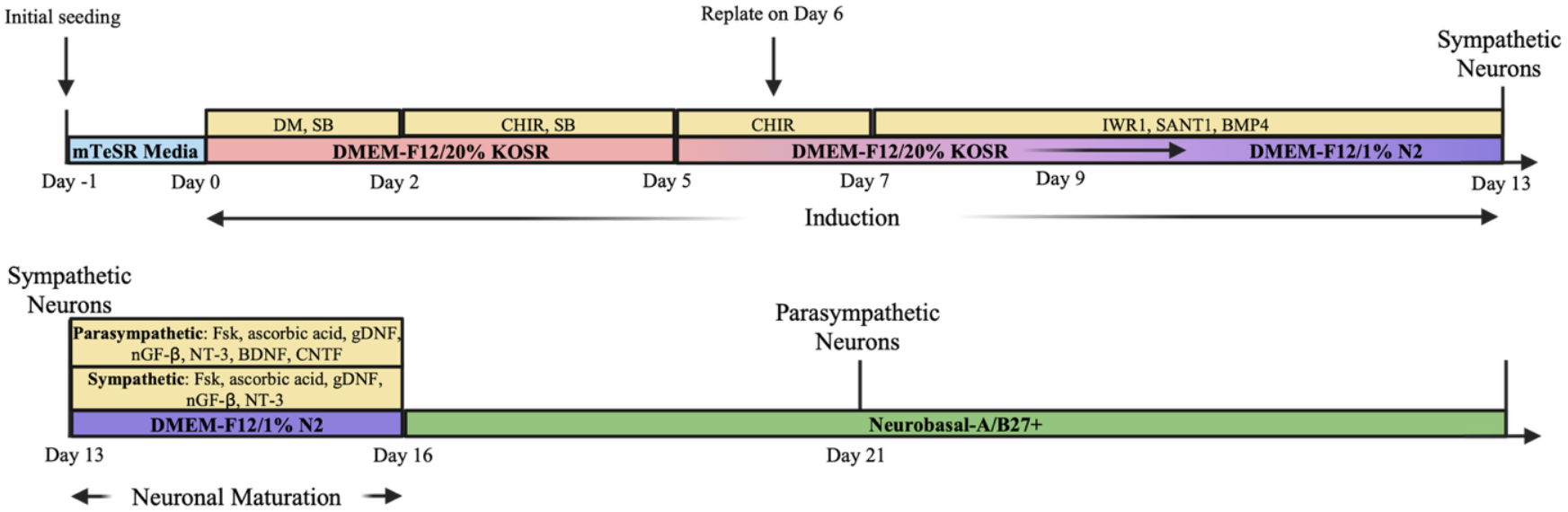
Differentiation protocol for sympathetic and parasympathetic motor neuron populations.

### HAEC Cultures

Human primary aortic endothelial cells (HAEC) (ATCC PCS-100-011, Lot: 80923231) were maintained in BBE Endothelial Cell medium (Vascular Cell Basal Medium, ATCC PCS-100-030 with Bovine Brain Extract (BBE) Endothelial Cell Growth Supplement, ATCC PCS-100-040) at standard cell culture conditions (5% CO_2_, 37°C) in cell culture flasks. When cultures reached confluency, cells were dissociated with 0.25% v/v trypsin-EDTA (Gibco 25200056) and seeded at densities between 2,500 to 7,500 cells/cm^2^.

### Neurovascular MPS Seeding

GelMA was synthesized as previously published, using fish gelatin^18^. GelMA was diluted to 5% w/v in a Day 5-7 medium solution with 0.5% w/v LAP. Day 6 NPC derived from human iPSC and P6-P7 HAMSC were each resuspended in 5% GelMA precursor solution, each at 5 x 10^6^ cells per mL. 27 μL of HASMC/GelMA precusor solution were seeded into the middle chamber of the organ chip and crosslinked for 45 seconds with blue light (405 nm, 10 W), with the glass side of the chip facing the lamp. 27 μL of NPC/GelMA precusor solution were seeded into each outer chamber of the organ chip and crosslinked for 130 seconds with blue light, with the glass side of the chip facing the lamp. Medium was exchanged at ∼20 min after seeding to remove any uncrosslinked GelMA precursor solution. On Day 5 of culture P5 HAEC were seeded at 5 x 10^5^ cells/cm^2^ onto the middle 2D chamber.

### Disassembly

For disassembly, first the screws and nuts were removed from each MPS. The layers were then loosened by gently swiping a screwdriver along the edges of the assembly. The media reservoir and 2D layer was separated from the 3D layer by twisting the pieces away from each other and pulling them apart. The 2D layer was then gently pulled apart from the media reservoir (Fig. 6).

**Figure 6.**
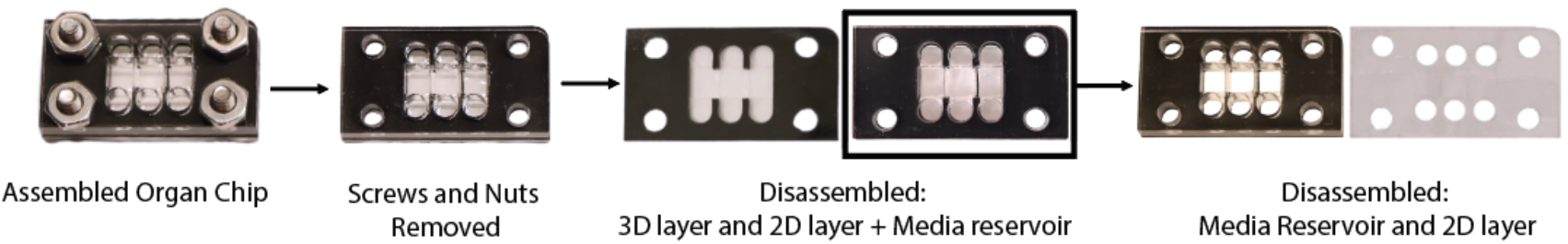
Demountable chambers support disassembly at culture endpoints.

### Immunostaining

Following disassembly of the organ chip, 3D layers of the organ chip were placed in a 6-well plate. Samples were rinsed with HBSS. Samples were then fixed in 4% paraformaldehyde (ThermoScientific, 043368.9M) in HBSS (Gibco 14065-056) for 40 min and permeabilized in 0.1% Triton X-100 (ThermoScientific A16046.AP) in HBSS for 30 min at room temperature, each followed by three 10-minute washes in HBSS. The samples were then blocked by 2.5% goat serum in HBSS (blocking solution) overnight (∼16 hours), followed by an overnight (∼16 hours) incubation with primary antibodies diluted in the blocking solution, both at 4°C on a rocker. Following four 1-hour HBSS washes on a rocker to remove excess primary antibody solution, the samples were incubated with secondary antibodies diluted in blocking solution overnight (∼16 hours) at 4°C on a rocker. Following four 1-hour HBSS washes on a rocker to remove excess secondary antibody solution, samples were incubated with DAPI (Invitrogen D1306) diluted 1:1000 in blocking solution for 10 minutes. Finally, three 10-minute HBSS washes were performed on a rocker to remove excess DAPI solution.

The 2D layer was separated from the assembly layer (Fig. 6). The edges of the layer were cut, and samples were placed into a 12-well plate. Following a rinse with HBSS, samples were fixed in 4% paraformaldehyde (ThermoScientific, 043368.9M) in HBSS (Gibco 14065-056) for 10 min and permeabilized in 0.1% Triton X-100 (ThermoScientific A16046.AP) in HBSS for 10 min at room temperature, each followed by three 10-minute washes in HBSS. The samples were then blocked by 2.5% goat serum in HBSS (blocking solution) overnight (∼16 hours), followed by a 2-hour incubation with primary antibodies diluted in the blocking solution at room temperature. Following three 5-minute HBSS washes on a rocker to remove excess primary antibody solution, the samples were incubated with secondary antibodies diluted in blocking solution for 1 hour at room temperature. Samples were washed three times for 5 minutes each with HBSS to remove excess secondary antibody solution. Samples were mounted with Prolong DAPI and a #1 thickness 18 mm x 18 mm coverslip. Samples were then sealed using nail polish.

All samples were imaged using an inverted fluorescence microscope (Zeiss Axio Observer). Images were processed using ImageJ/FIJI software.

### Computational Fluid Simulations

The laminar flow of cell culture medium through the organ chip flow component was modeled in COMSOL. First, SolidWorks was used to create the 3D flow channel, which was then exported as an STL file and imported into COMSOL as the geometry. Fluid properties of water were used, with the viscosity manually changed to 9.35 x 10^-4^ Pa-S, averaging previously reported values^19^, to more closely model the viscosity of DMEM cell culture medium. Using the laminar flow module, no slip walls and fully developed flow were selected, and a volumetric flowrate of 8 x 10^-8^ m^3^/s was set at the inlet (equivalent to 4.8 mL/min, a flowrate compatible with most peristaltic pumps which are frequently used in vascular studies). Backflow was not suppressed at the outlet to model any possible recirculating fluid. A stationary study was run with meshing set to “normal” and then the shear stress was modeled by the input “spf.sr * spf.mu” to multiply the shear rate and dynamic viscosity.

## Results

### Elastomer Selection and Impact on Viability

A materials selection matrix weighted for stability, gas permeability, cost, machineability (laser cutting), and optical transmission yielded four commercially available elastomers. Small samples were added directly to F11 2D cultures on well plates and evaluated for toxicity. Silicone, neoprene, and Viton did not impact viability compared to controls (Fig. 2), while EPDM negatively impacted F11 cell health. Viton was ultimately selected for is inertness, low gas permeability, and ease of manufacturing (Table 1).

### MPS with Removable Layers

In our design, a combination of cut-and-assemble fabrication and SLA 3D printed components were successfully utilized to construct a 3D neurovascular MPS, allowing us to construct precise geometries with tunability in the X, Y, and Z axes. SLA was also used to fabricate the example flow component (Figs. 3 & 9), reducing the need for multiple tape layers, which therefore also reduces the possibility of leaks from loosening of the assembly. Additionally, our design avoids any contact of cells directly with the 3D printed components, thus reducing the importance of resin biocompatibility, as the cells do not directly contact the 3D printed coverslip holder or flow components.

The neurovascular MPS includes three internal structures: a 3D cell culture layer, a 2D cell culture layer, and a media layer (Figs. 3 & 4). The 3D cell culture layer allows the contact of adjacent hydrogel culture chambers through GelPins^17^, forming a contiguous culture, and contains a glass bottom to facilitate high resolution microscopy. In the 2D culture layer, cells are seeded on a semi-permeable membrane, allowing contact with the hydrogel-encapsulated cells below. For assembly, the three layers are stacked with a 3D printed piece at the bottom of the assembly and held together using stainless steel screws and nuts. The three layers of the organ chip were easily removed from each other by disassembling the hardware and gently pulling the layers apart (Fig. 6), allowing direct access to each individual cell type (Fig. 7).

**Figure 7.**
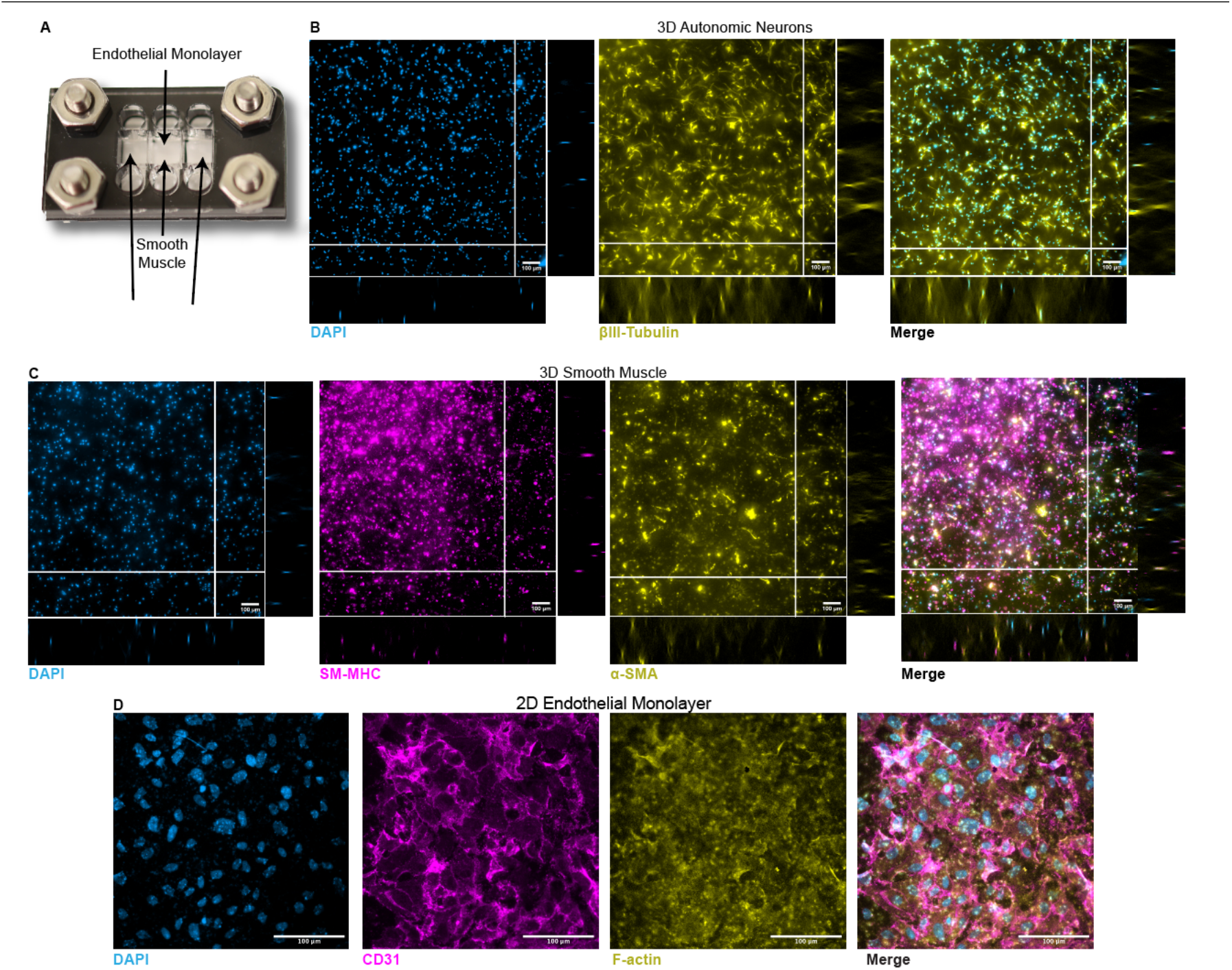
Demountable platform facilitates endothelial, smooth muscle, and neuron cultures for up to 27 days on chip. A) Seeding locations on chip. B) Differentiated neurons exhibit an adherent morphology with budding neurites within 3D GelMA chamber. Maximum intensity projection of slices 1-95 of ANS neuron populations. D) 3D smooth muscle culture exhibits heavy chain myosin, alpha smooth muscle actin, and expanded morphology. Maximum intensity projection is shown. D) 2D endothelial monolayer. (Scale Bars = 100 μm)

### Immunostaining and visualization of neural outgrowth in 3D culture

Organ chips with sympathetic or parasympathetic neurons were disassembled on Day 19 or Day 27 of neuron differentiation, and the 3D culture layer was set aside for imaging (Fig. 7B). The 3D culture layers were then fixed, permeabilized, and immunostained. Samples were imaged using an inverted fluorescence microscope (10x objective) to visualize expression of β-III tubulin (B3T), a neuronal marker, and cell nuclei (counterstained with DAPI). Z-stack images were taken, and maximum intensity projections of slices 1-95 of Day 27 parasympathetic neurons (Fig. 7). As expected, expression of β-III tubulin was present and the cells were distributed throughout the height of the hydrogel, as observed in the orthogonal views, demonstrating innervation of the chamber. Therefore, using our nerve-artery organ chip platform, we were able to demonstrate the visualization of expression and distribution of neuronal markers in differentiating neural cells on chip.

After immunostaining, organ chips were imaged using a 63x oil immersion objective (Fig. 8). For immunostaining, organ chips with sympathetic or parasympathetic neurons were dissembled on Day 19 or Day 27 of neuron differentiation, and the 3D culture layer was set aside for imaging. The 3D culture layer was then fixed, permeabilized, and immunostained for β-III tubulin (B3T - neuronal marker) and tyrosine hydroxylase (TH – sympathetic neuron marker; Fig 8A) or choline acetyltransferase (ChAT – parasympathetic neuron marker; Fig 8B) for organ chips with sympathetic or parasympathetic neurons, respectively, and cell nuclei were counterstained with DAPI. As the 3D culture layer of the chip can be easily maneuvered after removal from the overall chip assembly, we were able to image the chip easily with a 63x oil immersion lens. Qualitatively, there is presence of TH in the sympathetic differentiation group on Days 19 and 27, while there is an increased presence of ChAT on Day 27 of the parasympathetic differentiation group compared to Day 19.

**Figure 8.**
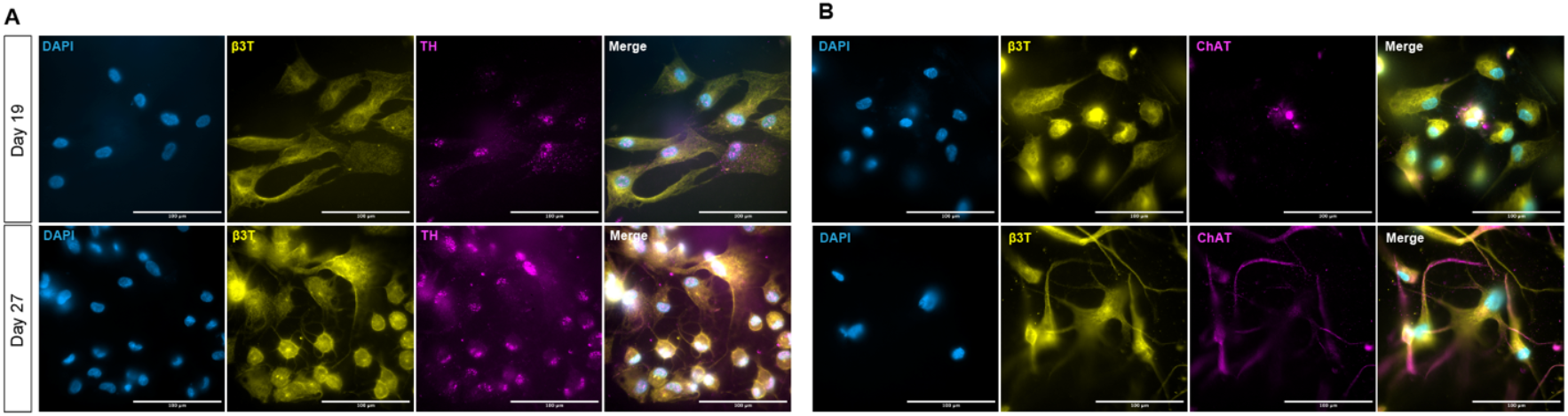
Demountable chambers support high resolution imaging of 3D tissue chambers, 63x oil immersion, on a standard inverted fluorescent microscope., as demonstrated by staining of neural and A) sympathetic markers and B) parasympathetic markers. (Scale Bars = 100 μm)

### Computational fluid simulations demonstrate high and low shear stress arterial flow on chip

Computational fluid simulations were run in COMSOL software to demonstrate flow on chip (Figs. 3C and 9). Flow channels were designed in SolidWorks and exported to COMSOL for flow simulations. Use of CAD allowed rapid redesigning of the flow channels, and this was utilized to achieve a range of physiologically relevant flow rates and shear stress by varying the height of the flow channel while preserving the rest of the geometry. A channel height of 0.25 mm was used for the high shear stress flow simulation, while a channel height of 0.5 mm was used for the low flow simulation. In the EC region (roughly outlined in black), a shear stress of ∼1.1 Pa and 0.3 Pa was achieved over the endothelial region to model high and low shear stress in arteries. To demonstrate potential for application in benchtop experiments, an example flow attachment was 3D printed by SLA and attached to the chip, and distilled water with blue food coloring was flown through by a syringe pump for visualization (Fig. 3C). Direct comparison of the high vs. low shear stress (Fig. 9).

**Figure 9.**
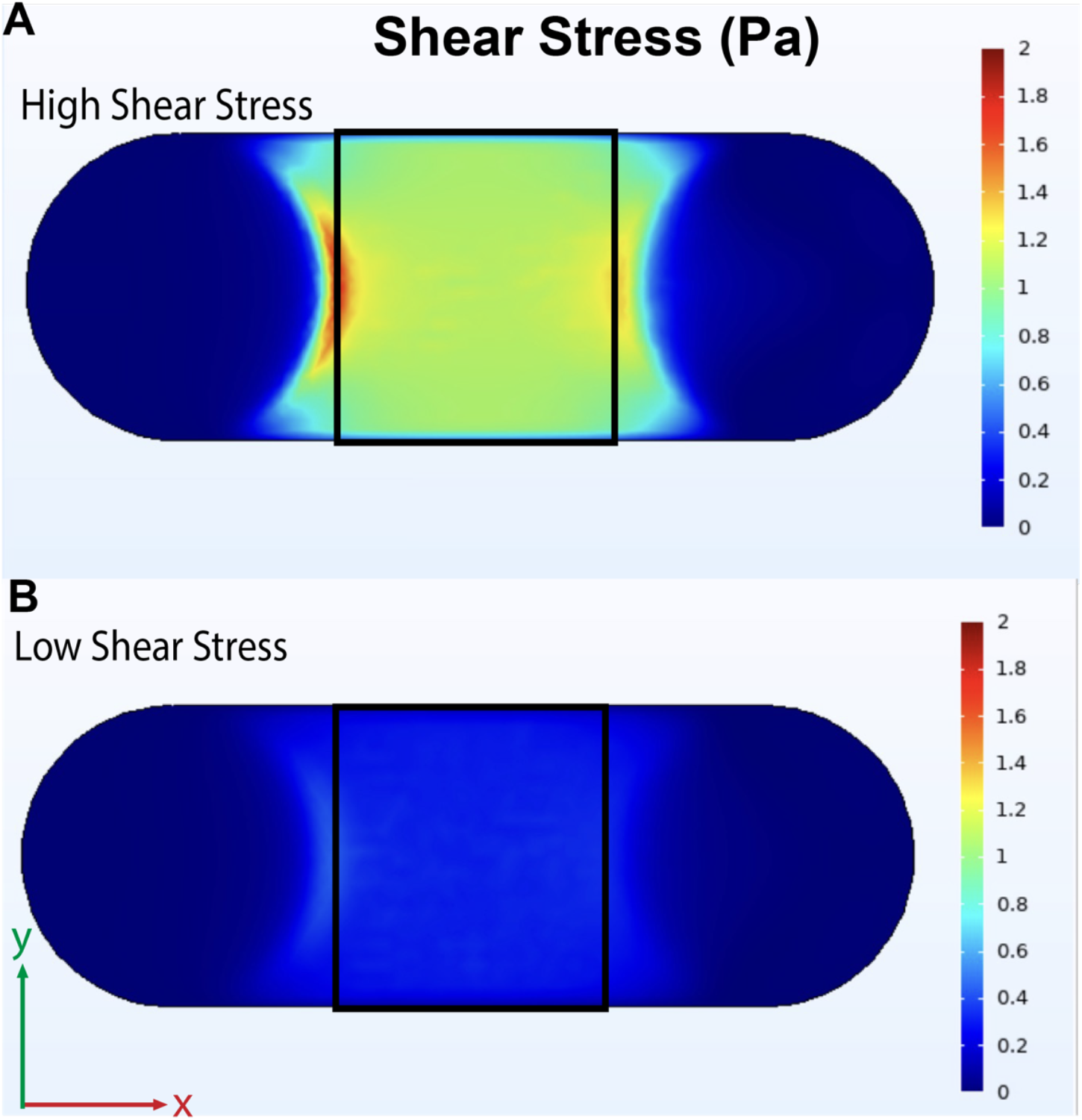
High and low shear stress arterial flow on chip. The view from the cellular region is shown, with the inlet on the left side. A) High and B) low shear stress (bottom) flow regimes were achieved on chip by varying the flow channel height.

## Discussion

Future translation of MPS technologies relies on approaching the complexity and heterogeneity of *in vivo* tissue yet retaining ease of use, including capabilities for traditional microscopy techniques without the need for expensive specialized equipment^20,21^. Our layer-by-layer approach to MPS assembly allows limitless fabrication of elements or chambers in the z-direction. While beneficial for increasing the complexity and bio-relevancy of these platforms, this engineering advantage is at the determent of traditional optical microscopy techniques in irreversibly sealed devices. Therefore, demountable platforms allow high resolution imaging with standard inverted fluorescent microscopes equipped with standard working distance objectives.

The removable interfacing 2D and 3D cell cultures on our organ chip provide several advantages, from high resolution endpoint analyses to inclusion of flow components. In addition to the direct advantages of higher imaging resolution and accessibility for endpoint assays, separation of 2D and 3D layers allow for optimal incubation times for specific live and/or fixed assays on each layer. Compared to a sealed 3D system, the demountable chambers also enable tissue-specific immunocytochemical staining for each chamber to allow for each filter cube/laser setting/secondary antibody to be utilized for each MPS section, providing expanded resolution of standard fluorescent microscopy. Several studies have employed multiplexed imaging for investigations in oncology research^22-24^. However, only a few studies exploring multiparametric imaging on organ chips^20,25^, and reconfigurable or reversibly sealed systems, such as the one we present here, can facilitate further integration of multiplexed approaches on organ chips. Overall, the removable, interfacing chambers of our organ chip can be easily redesigned, expanded, and employed for understanding developmental mechanisms and disease pathophysiology, especially by employing multiple differentiating human cell types derived from stem cells.

The use of widely available CAD and CFD software as well as common makerspace equipment reduces time and cost of organ chip fabrication and redesigns. Because of these advantages, use of 3D printed parts for organ chips has recently been explored. Biocompatibility is a primary concern with using 3D printed resins for cell culture, and there has been some investigation on material composition, resolution, and post-print processing for use in cell culture devices^26-29^.Particularly, there have been several reports of the use of FormLabs Clear Resin, a methacrylate-based resin which we used here. Hart et al. reports HL-1 (immortalized rat cardiomyocyte) viability of greater than 96% with FormLabs Clear Resin following a 70% isopropanol wash and a UV post-cure^30^. However, others report lower viability and/or cell attachment to FormLabs clear resin after isopropanol washes and a UV post-cure ^31,32^, including no significant growth of human umbilical vein endothelial cells growth after 24 hours ^33^. As our organ chip design avoids direct contact of cells with the FormLabs clear resin, possible biocompatibility concerns are mitigated. 3D printing may contribute to bridging the gap to widespread adoption of organ-on-chip, though further standardized studies on biocompatibility and other designs avoiding contact of 3D printed parts with cellular components must be explored.

Manually re-sealing the chip may cause slight changes in alignment and/or non-uniform load, and hardware components are prone to mechanical wear and tear, as previously discussed^11^. Affordable torque screwdrivers ($50-$100) are valuable tools for dialing the proper torque to eliminate the risk of leaking, why not overtightening assembly. For our hardware, a torque of 0.12 Nm met this need and could be easily checked over the duration experiment since thermal cycles of normal cell culture techniques was found to influence fastening torque.

Cell culture inserts offer one solution for demountable devices^15,34-38^, some of which include commercially available Transwell® cell culture inserts^39-49^. Transwell® inserts are a standard approach for membrane-separated systems, and their inclusion in organ chip devices enhances reproducibility and also allows the addition of flow^39^. Several cell culture insert based organ chips have components which can be autoclaved and/or reused ^41-43,46^, and several allow increased throughput^37,42,43^, which may both improve reproducibility and reduce the time needed for repeated device fabrication. However, these systems are limited in their ability to recapitulate 3D tissues and imparting controllable flow regimes to apply physiologically relevant shear stress. Further development of organ chip systems with rapidly produced or commercially available components, especially those that incorporate 3D cell cultures, may improve accessibility and utility of organ chip platforms for mechanistic studies.

As suggested by Teixeira Carvalho et al., our organ chip provided a physiologically relevant environment for stem cell differentiation by controlling the spatial orientation of cell types^11^. Additionally, all cell culture chambers can be directly accessed for endpoint assays, which is often not possible in permanently sealed systems. This feature can be further utilized to extract the cells from the organ chip, especially by using hydrogels compatible for such assays. For example, RNA extraction methods have been reported from several clamped organ chips^11^. Some methods of protein extraction have also been reported for hydrogel encapsulated cells but are comparatively limited^34,50^. Application and further development of these methods for the organ chip could be very useful for mechanistic investigations of multiple cell types in co-culture.

Though we demonstrated a neurovascular culture in this work, the applicability of the organ chip presented here is not limited to the nerve-artery interface. Various cell types can be cultured to model other vascularized or innervated systems, or other systems altogether. Additionally, a membrane with larger pores could be employed to study cell migration in various environments, such as cancer cell migration, angiogenesis, or immune cell migration, in multi-tissue systems. Overall, the unique capabilities of this platform may be useful for mechanistic studies of complex multi-tissue systems.

## Acknowledgements

The authors would like to thank Dr. Max Winkelman for preliminary work that led to the development of this work, and Nolan Burson for assistance with cell culture. Also, the authors would like to thank Dr. Richard West’s Lab at Northeastern for COMSOL support.

## Author Contributions

SB, AK, GD, and RK conceived the project. SB, RPB, and LA performed all experimental work and wrote the manuscript. SB and RPB designed and fabricated organ chips, and BS designed and optimized fabrication of stereolithography printed components. SB and DM performed image analysis, with support from AK and RK. AK, RK, and GD provided intellectual feedback and support. All authors edited and provided feedback on the manuscript. RK supervised the work.

## Additional Information

Competing financial interests: The authors declare no competing interests.

